# Necrotizing enterocolitis-induced systemic immune suppression in neonatal preterm pigs

**DOI:** 10.1101/2020.09.21.306290

**Authors:** Shuqiang Ren, Xiaoyu Pan, Yan Hui, Witold Kot, Fei Gao, Per T. Sangild, Duc Ninh Nguyen

## Abstract

**Objectives:** Preterm infants are at high risks of sepsis and necrotising enterocolitis (NEC). Some develop sepsis shortly after suspected or confirmed NEC, implying that NEC may predispose to sepsis but the underlying mechanisms are unknown. Using NEC-sensitive preterm pigs as models, we investigated the immune status in animals with and without NEC.

**Methods:** Preterm pigs (n=113, caesarean delivered at day 106) were reared until day 5 or 9. Blood was analyzed for T cell subsets, neutrophil phagocytosis, transcriptomics and immune responses to LPS challenge. Gut tissues were used for histology and cytokine analyses. Pigs with/without macroscopic NEC lesions were scored as healthy, mild or severe NEC.

**Results:** Overall NEC incidence was similar on days 5 and 9 (61-62%) with less severe lesions on day 9, implying gradual mucosal repair following the early phase of NEC on day 5. Pigs with NEC, especially severe NEC, showed decreased goblet cell density and increased MPO^+^ and CD3^+^ cell density in the distal intestine or colon. Circulating parameters were minimally affected by NEC on day 5, but widely altered on day 9 in pigs with NEC, especially severe NEC, to the direction of immune suppression. These included elevated Treg frequency, impaired neutrophil phagocytosis, diminished LPS-induced cytokine secretions and immune gene responses, and consistently low expressions of genes related to innate immune signalling and Th1 polarization.

**Conclusion:** We shows evidence for NEC-induced systemic immune suppression, even with mild and sub-clinical NEC lesions, thereby suggesting mechanisms for increased secondary infections in infants with previous NEC diagnosis.

## Introduction

Preterm birth (before the completion of 37 weeks of gestation) occurs for approximately 10% of total pregnancies worldwide and is responsible for multiple early life complications and one million deaths every year^1^. Preterm infants are particularly susceptible to systemic infection, late-onset sepsis (LOS) and gastrointestinal diseases, including necrotizing enterocolitis (NEC)^1,2^. For many decades, these infectious diseases have been speculated to be related to the immature intestinal and systemic immune system with poor capacity to mount proper response to exogenous challenges, including gut bacterial colonization and enteral feeding^3–6^. Recently, it has been evident that the preterm newborn immune system is programed to a status of disease tolerance with immune suppression and minimized glycolytic activity^7,8^, which prioritizes cellular energy for maintenance of organ functions rather than hyper-inflammatory responses^9^. However, when a tolerable threshold of the immature gut and circulation is exceeded, elevated gut and systemic inflammation may occur via mucosal and systemic immune activation in an effort to resolve excessive challenges^6^. This hyper-inflammatory status is typical shortly before and at the diagnostic phase of NEC and LOS in preterm infants^6,10^.

The association of NEC and sepsis with increased systemic inflammation has led to multiple explorations of diagnostic biomarkers for these diseases^11^. Previous studies in human NEC patients and animal models have shown that NEC progression is associated with the elevated fraction of systemic IL-17 producing Treg ^12^ and increased levels of acute phase molecules (SAA, IL-6, IL-8, CRP)^13^, as well as diminishment of systemic anti-inflammatory molecules (e.g. TGF-ß2)^14^. Similar host responses are also observed for LOS^15^. Of note, septic neonatal and elderly patients gradually develop immunosuppression after diagnosis, which predispose to increased risks of secondary infection and organ failures^16,17^. However, it is unknown whether NEC patients following surgery or medical treatment also possess an immune suppressive status that may predispose them to secondary infection and sepsis.

At clinical diagnosis of NEC, antibiotics are indispensably treated to decrease gut bacterial overgrowth and translocation^18,19^. Some of the NEC patients recovering from medical treatments or surgery later develop LOS, suggesting bacterial translocation from the compromised gut barrier to the circulation preceding LOS^20,21^. However, it is also possible that NEC lesions or the interventions associated with NEC including antibiotics alters the systemic immune system, thereby together with gut bacterial translocation predisposing to LOS. Long-term usage of antibiotics are well-known to suppress immune functions and increase infection risks^22,23^. However, it remains elusive if NEC or antibiotic treatment at clinical NEC diagnosis can separately induce immune suppression and predispose to secondary infection and sepsis.

Investigation of the isolated effects of NEC on immune status is not possible in preterm infants as it is considered unethical not to treat NEC patients with antibiotics. Alternatively, the preterm pig is well-acknowledged as a clinically relevant model of NEC because it assimilates most complications in preterm infants including impaired mucosal and systemic immune system, immature organ functions and spontaneous NEC development following formula feeding^3–5,24^. In addition, it is possible to induce sub-clinical NEC lesions with various severities in formula-fed preterm pigs and further rear them without antibiotic treatment^25^, in order to investigate the isolate NEC effects on organ systems. Based on this background, we hypothesized in the current study that NEC lesions induce systemic immune suppression in preterm neonates, thereby impairing immune competence against secondary infectious challenges. To test the hypothesis, we reared preterm pigs with bovine milk diets to induce sub-clinical NEC at various degrees of severity (mild or severe, based on macroscopic lesions) and investigate their systemic immune status.

## Results

### Gut inflammation associated with NEC

With similar feeding regimes across ten litters, 36/113 pigs were planned for euthanasia on d 5 and the remaining pigs were reared until d 9. All pigs in the cohort survived until the planned euthanasia without severe clinical symptoms. Sub-clinical NEC, diagnosed from the macroscopic scoring at euthanasia, appeared in 22/36 pigs (61%) on day 5 and 48/77 pigs (62%) on d 9 without statistical significance (P = 0.53, Fig. 1A-B). In contrast, the incidence of severe NEC was reduced on d 9, relative to d 5 (16/77, 21% vs. 15/36, 42%, P < 0.05, Fig. 1B). This suggests that NEC lesions were already induced on d 5 in these pigs and partly healed in the following days. This phenomenon may be similar to that in formula-fed preterm infants with possible sub-clinical lesions gradually being healed without clinical symptoms^26,27^.

**Figure 1.**
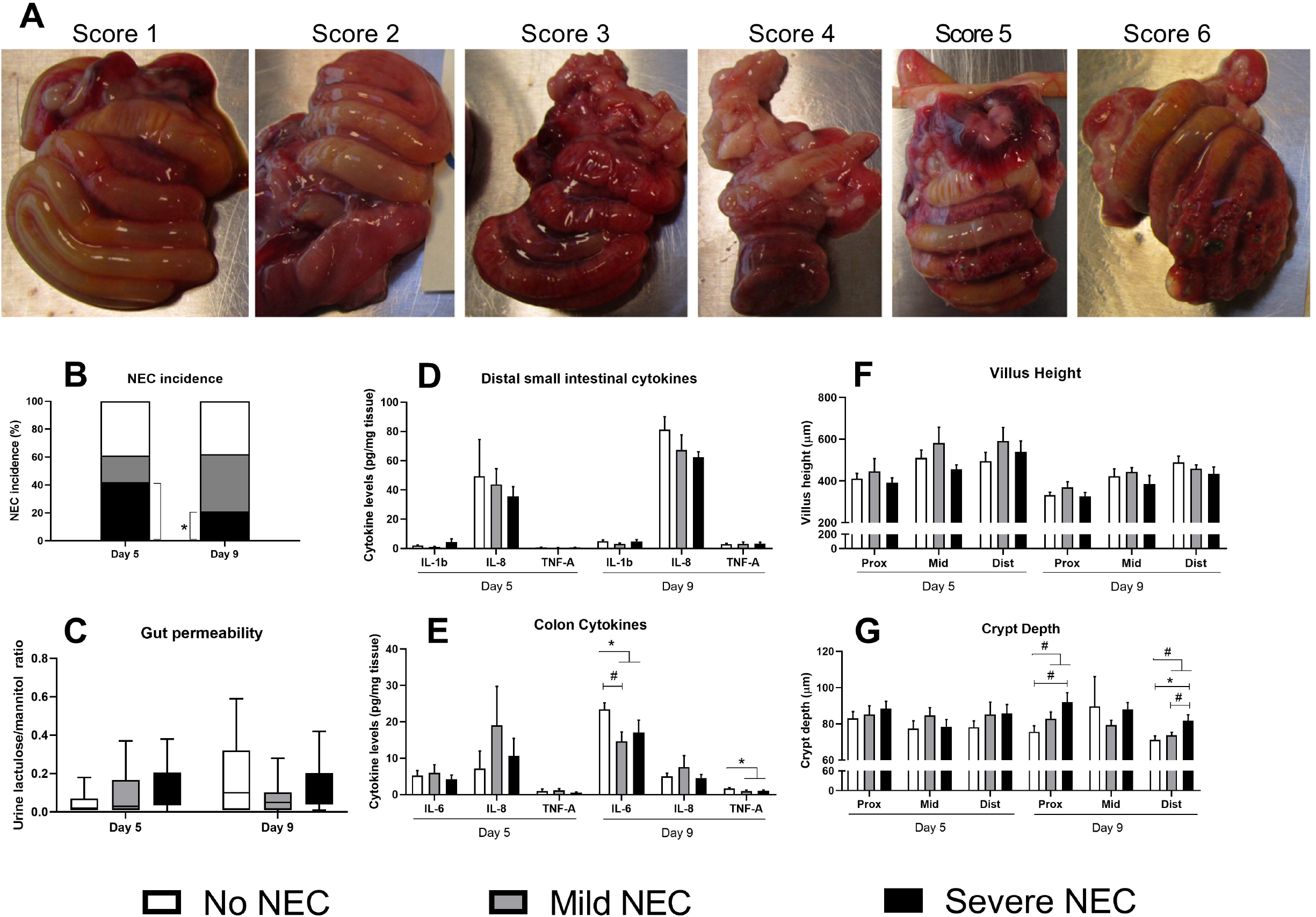
Incidence of NEC and related gut parameters. NEC score, overall incidence of NEC and incidence of mild and severe NEC (A-B). Gut permeability (C, n = 6-9 and 8-22 on d 5 and 9, respectively). Cytokines in the distal small intestine and colon (D-E, n=4-6 and 5-14 on d 5 and 9, respectively). Villus height and crypt depth (F-G, n = 7-15 and 9-27 on d 5 and 9, respectively). Values in B-F are means ± SEM. * and ** P < 0.05 and 0.01, respectively. # P < 0.1.

Pigs with mild and severe NEC showed numerically higher values of gut permeability than their healthy littermates on d 5, but not d 9 (Fig. 1C), supporting the data of less severe NEC after d 5. All tested inflammatory cytokines in the distal small intestine did not show any differences at any time points between pigs with vs. without NEC (Fig. 1D) but interestingly IL-6 and TNF-α levels in colon tissues on d 9 were lower in pigs with mild and severe NEC, relative to those without NEC (P < 0.05, Fig. 1E). Gut morphology via H&E staining showed no differences in villus height at both time points (Fig. 1F), but crypt depth values were higher in pigs with NEC, especially severe NEC, vs. without NEC (P < 0.05, Fig. 1G).

Histological data revealed a drop in mucin-containing goblet cell density in the colon of pigs with NEC (P < 0.01), especially severe NEC (P < 0.05), on d 5 but not d 9 (Fig. 2A-D). No difference in goblet cell density was detected in the distal small intestine. On d 9, the density of MPO-positive cells (neutrophils/macrophages) tended to be higher in the distal small intestine of pigs with vs. without NEC (P = 0.1), while it was much higher in the colon of pigs with severe vs. mild or no NEC (P < 0.01, Fig. 2F-H). For the intensity of CD3-positive cells (T lymphocytes) on d 9, there was no difference in the distal small intestine but higher values in pigs with severe vs. mild NEC in the colon (P < 0.05, Fig. 2I-L). No differences in levels of MPO- or CD3-positive cells in any of the investigated regions among the groups were found on d 5 (data not shown).

**Figure 2.**
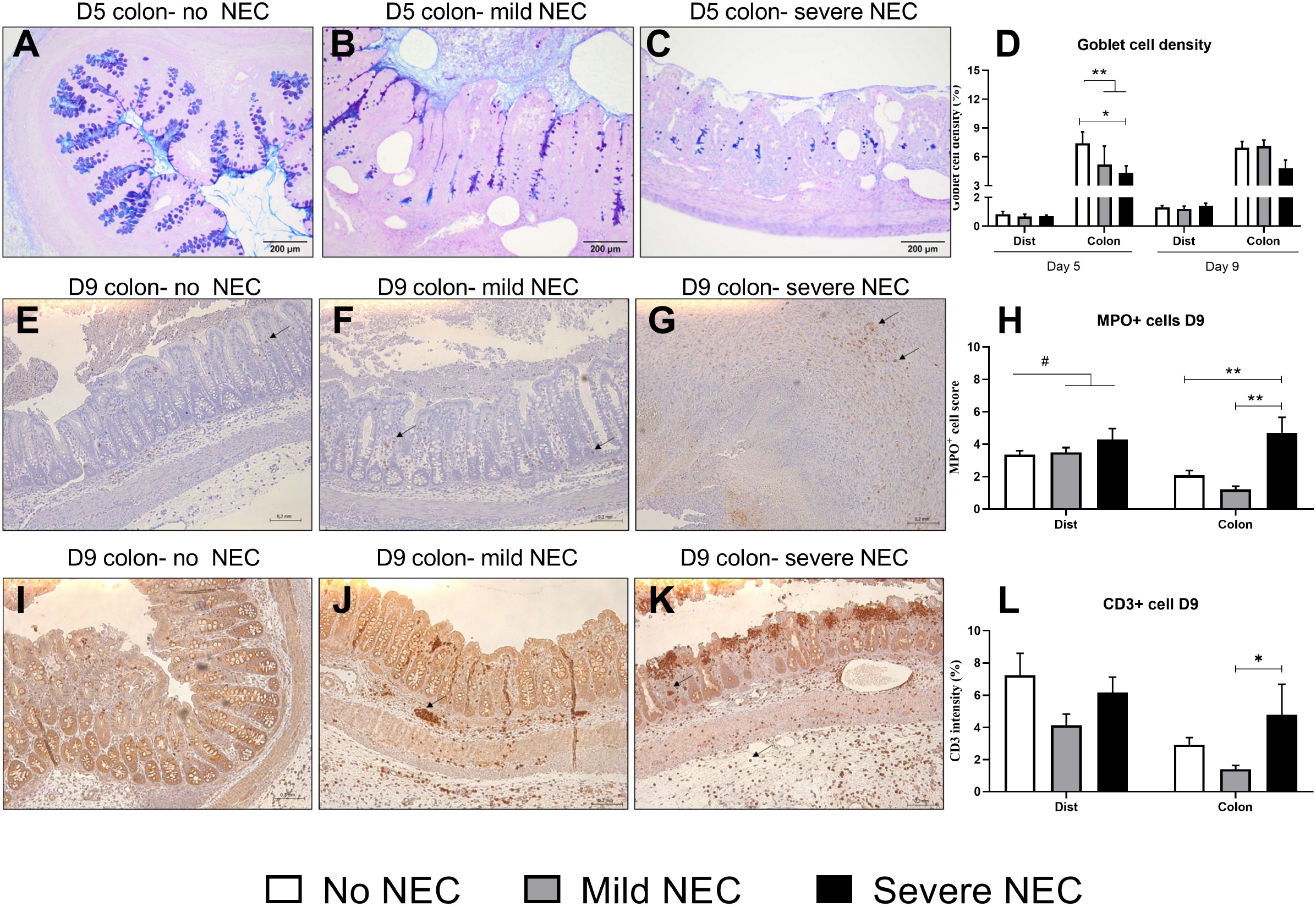
Gut histology in the distal small intestine and colon. Goblet cell density (A-D, n = 7-15 and 16-32 on d 5 and 9, respectively). MPO-positive cells (E-H, n = 4-14 and 5-14 on d 5 and 9, respectively). CD3 positive cells (I-L, n = 5-14 on d 5 and 9). Values in D, H, L are means ± SEM. *, ** P < 0.05 and 0.01, respectively. # P < 0.1. Lines in scale bars represent 200 μm.

### Gut microbiome

Gut microbiome, analyzed by 16S rRNA gene amplicon sequencing, was similar between pigs with vs. without NEC on both d 5 and 9, as assessed by Shannon alpha diversity (Fig. 3A), beta diversity with both unweighted and weighted Unifrac dissimilarities (Fig. 3B-C). Taxonomic comparison using ANCOM analysis showed dominant bacteria belonging to Enterobacteriaceae family on d 5 and dominant *Enterococcus* spp. on d 9 without any differences between pigs with vs. without NEC (Fig. 3D). This is in agreement with previous reports demonstrating no major significant association of gut microbiota alteration with NEC in human NEC patients during the first 4 weeks of life^28^.

**Figure 3.**
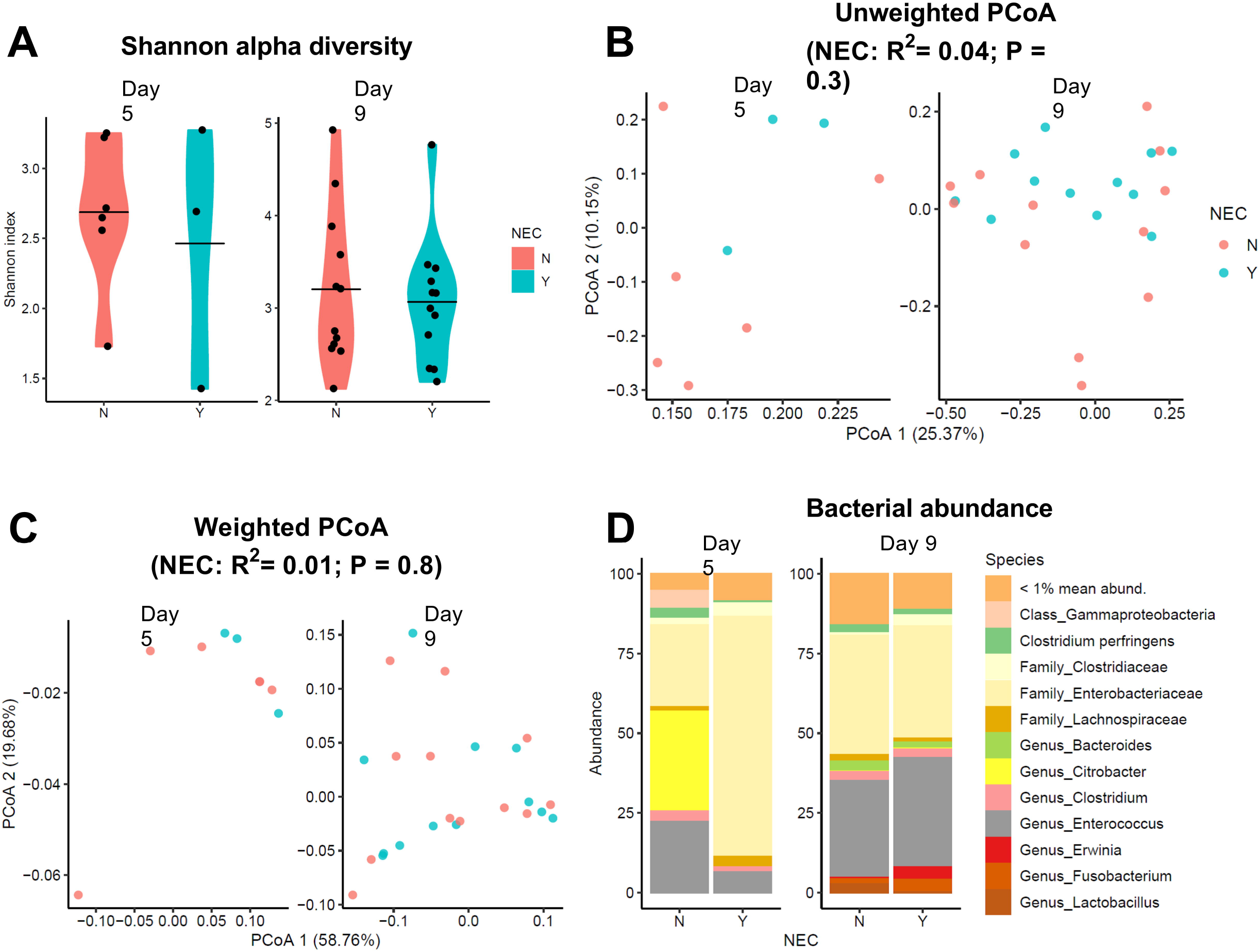
Colonic microbiome via 16S rRNA gene amplicon sequencing. (A) Shannon alpha diversity. (B-C) PCoA based on unweighted and weighted Unifrac distance metrics. (D) Relative bacterial abundance. n = 3-5 on d 5 and 12 on d 9. Labels with Y and N indicate pigs with and without NEC, respectively.

### Systemic immune status associated with NEC

To evaluate the systemic immune status associated with NEC at different time points, we compared pigs without vs. with mild vs. with severe NEC lesions with regards to their hematological profiles, various T cell subsets including blood regulatory T cells (Treg), neutrophil phagocytosis function, and cytokine secretion and gene expression in response to *ex vivo* whole blood challenge with LPS. Blood neutrophil counts on d 5 showed tendency to be higher in pigs with vs. without NEC (P = 0.06) and significantly higher in pigs with severe NEC (P < 0.05), but no difference among groups were detected later (d 9, Fig. 4A). Combined with data showing increased MPO-positive cells in the gut of pigs with NEC, especially severe NEC on d 9, the blood neutrophil data suggest more neutrophil production from the bone marrow at early phase of severe NEC and those cells gradually home to the gut at a later time point. Despite having similar neutrophil counts, pigs with severe NEC on d 9 showed lower number of neutrophils having phagocytic capacity than those without NEC (P < 0.05, Fig. 4B), indicating poorer innate immune functions at post-diagnosed NEC phase. Further, whole blood LPS challenge assay showed that TNF-α levels both before and after LPS challenge on d 9, but not d 5, was lower in pigs with vs. without NEC (P = 0.09 and < 0.05, respectively, Fig. 4C). No differences among groups were detected for IL-10 levels, except the increased levels after vs. before LPS stimulation (all P < 0.05, Fig. 4D). These data suggest systemic immune suppressive status or impaired Th1 response following NEC occurrence. This was supported by the data of immune suppressive cell subset Treg fraction (CD3^+^CD4^+^Foxp3^+^) showing no difference among groups on d 5 but significantly higher in those with vs. without NEC on d 9 (P< 0.01, Fig. 4E). These trends were also associated with lower levels of serum glucose in pigs with severe NEC (P < 0.05) only on d 9, but not d 5 (Fig. 4F). No or minor differences among groups were detected for the remaining hematologic parameters, fraction of T cells, helper T and cytotoxic T cells (Supplementary Table S4).

**Figure 4.**
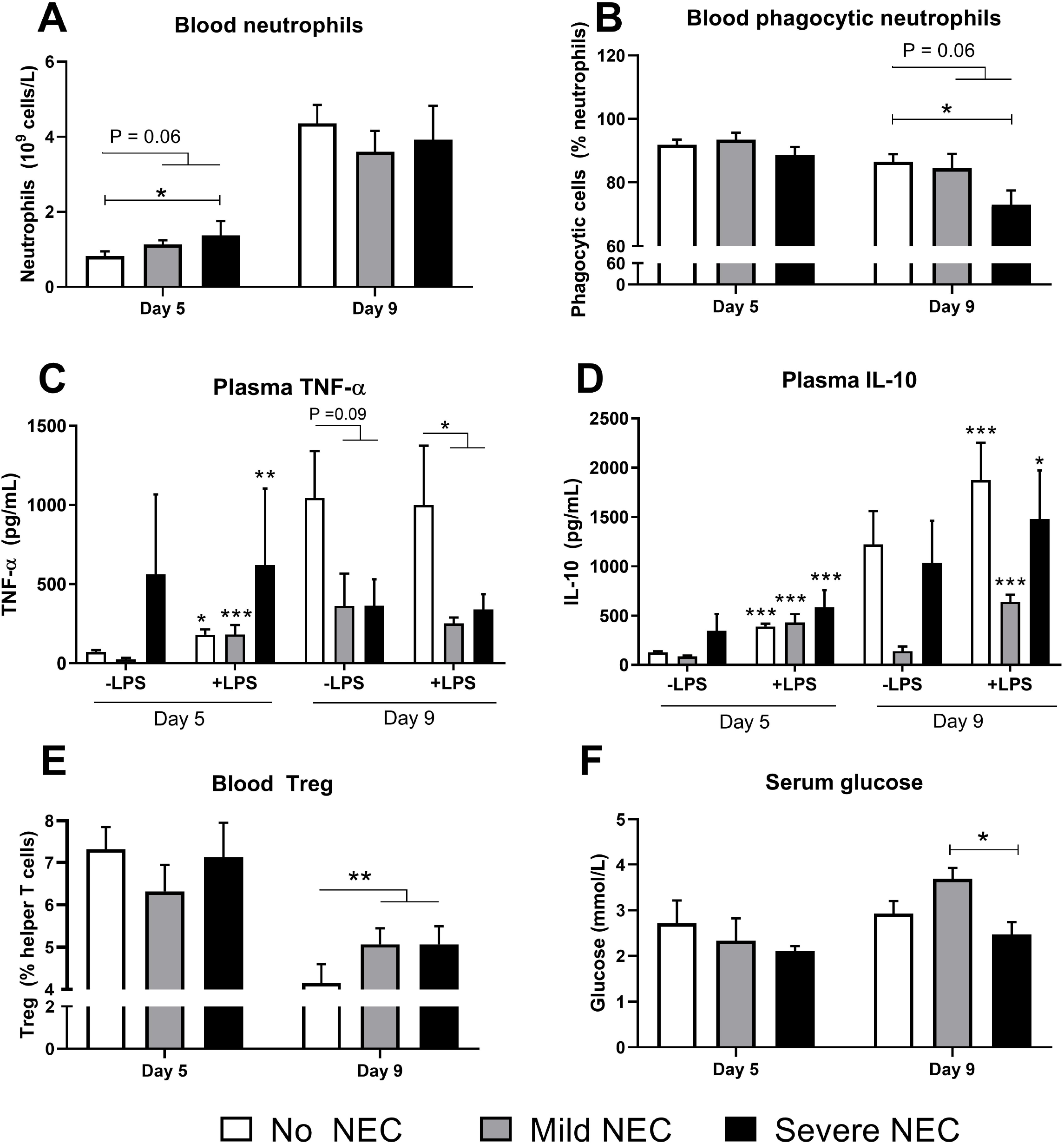
Systemic immune status associated with NEC. Blood neutrophils (A, n=13-27 and 14-21 on d 5 and 9). Blood neutrophil phagocytosis (B, n=5-14 and 4-10 on d 5 and 9). Plasma TNF-α and IL-10 without and with LPS stimulation (C-D, n= 5-13 and 4-9 on d 5 and 9). Blood regulatory T cells (Treg, E, n=15-26 and 11-16 on d 5 and 9). Serum glucose (F, n=3-10 and 9-28 on d 5 and 9). Values are means ± SEM. *, **, *** P < 0.05, 0.01 and 0.001, respectively.

We also examined a series of genes related to innate and adaptive immune responses following LPS challenge in whole blood on d 5 and 9 (Fig. 5 and Supplementary Table S5). On both d 5 and 9, pigs without NEC or mild NEC showed capacity to mount immune responses, when comparing LPS vs. no LPS stimulation (for 17/23 genes: *TLR2, TLR4, HIF1A, S100A9, TNFA, CXCL9, CXCL10, IL4, IL6, IL10, IFNG, TGFB1, HK1, PDHA1, PKM, RORC, PPARA*, P < 0.05 or 0.01 or 0.001, Fig. 5A-B, Supplementary Table S5). In contrast, pigs with severe NEC showed no responses to LPS stimulation for most of the investigated genes. When comparing gene expressions among the three groups (no NEC, mild or severe NEC) on d 5, most of the investigated genes showed no differences, except minor differences before LPS stimulation in *TLR2* and *S100A9* levels (Fig. 5A-B). Different from d 5, blood immunity gene expressions on d 9 consistently showed lower levels in pigs with NEC (either mild or severe or pooled mild and severe) vs. without NEC both before and after LPS stimulation. The trends applied to *TLR2, TLR4, S100A9, TNFA, IL4* and *IL10* before LPS stimulation, and *TLR2, HIF1A, S100A9, TNFA* and *IL10* after LPS stimulation (Fig. 5A-B). Collectively, both FACS and neutrophil functions, cytokine and qPCR data indicate increased immune suppression in pigs at post-diagnosed phase of NEC on d 9, but not d 5.

**Figure 5.**
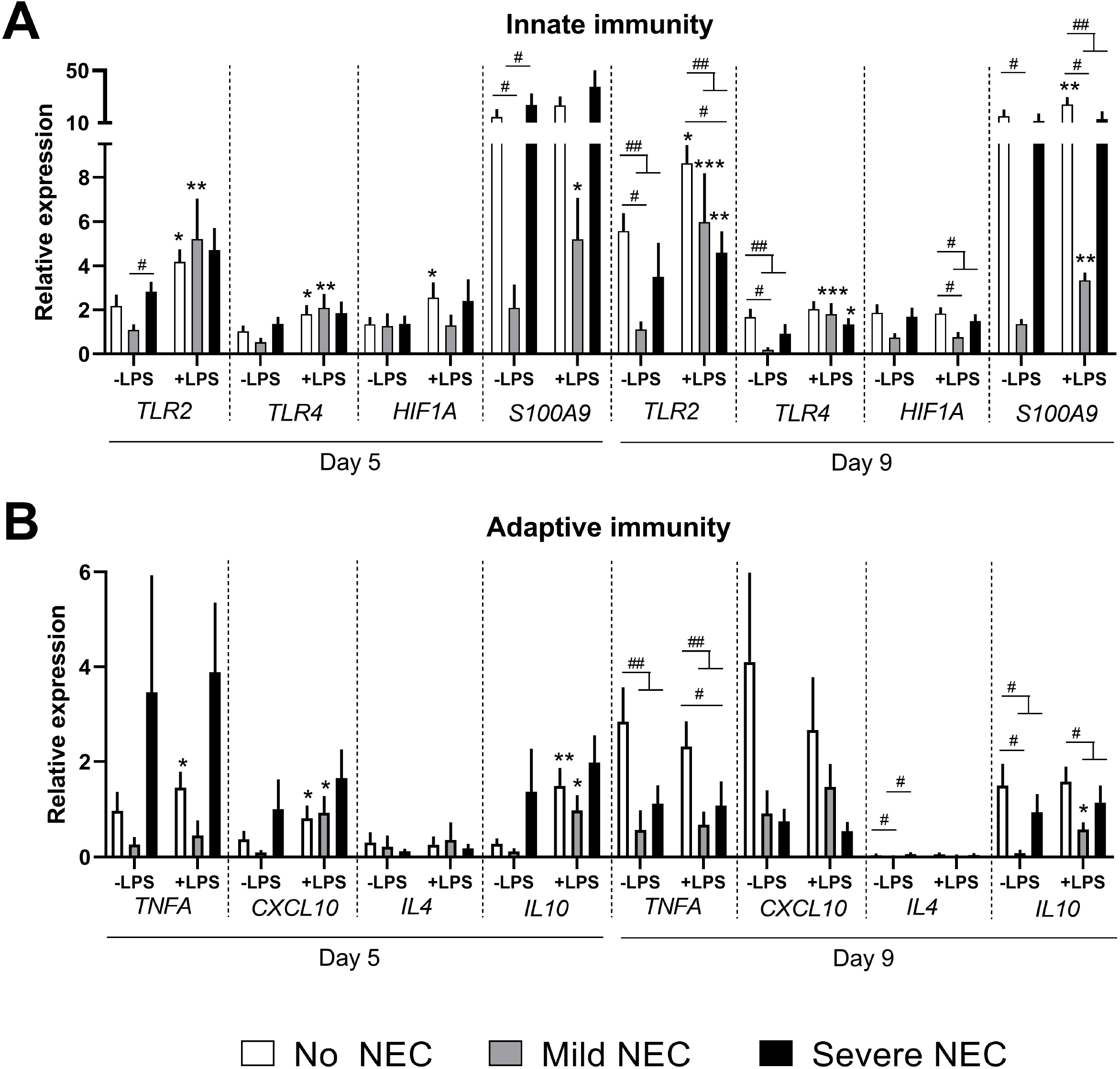
Blood gene expressions in LPS-stimulated whole blood of pigs with and without NEC. Innate (A) and adaptive (B) immunity-related genes. n=3-9 in each group on d 5 and 9. Values are means ± SEM. *, **, *** P < 0.05, 0.01 and 0.001, respectively, relative to the corresponding controls without LPS stimulation. #, ## P < 0.05 and 0.01 between the indicated groups.

### Blood transcriptomic profile associated with NEC

As differences in systemic parameters between pigs with vs. without NEC were mainly detected on d 9, we randomly selected 5 pigs with and 5 without NEC (matched control from the same litter) for blood transcriptomics to further profile possible immune-metabolic pathways associated with the immune suppressive status in NEC pigs. We identified 684 DEGs between pigs with vs. without NEC, with 378 down-regulated genes and 306 up-regulated genes in NEC pigs (Fig. 6A, Supplementary Table S1). Enrichment analyses demonstrated that NEC pigs showed down-regulated pathways related to both innate immunity and adaptive immunity (T cell receptor and JAK-STAT signaling, TNF signaling, chemokine signaling, Fig. 6B, Supplementary Table S2). Conversely, no pathways associated with up-regulated DEGs by NEC were enriched. Gene interaction network analysis among all DEGs showed top 10 key hub genes (based on betweenness) involved in TLR, IFN-gamma, neutrophil and T cell signaling (*CCR5, SOCS1, IGF2R, PIK3AP1*), energy production via oxidative phosphorylation (*ATP1A1, SOD2*) and 9 out of 10 hub genes were down-regulated in NEC pigs (Fig. 6C). Of note, NEC pigs also showed up-regulation of a series of genes associated with immune suppression, e.g. *TGFA, TGFBI, SMAD7*, involved in TGF-ß signaling (Fig. 6D, Supplementary Table S1).

**Figure 6.**
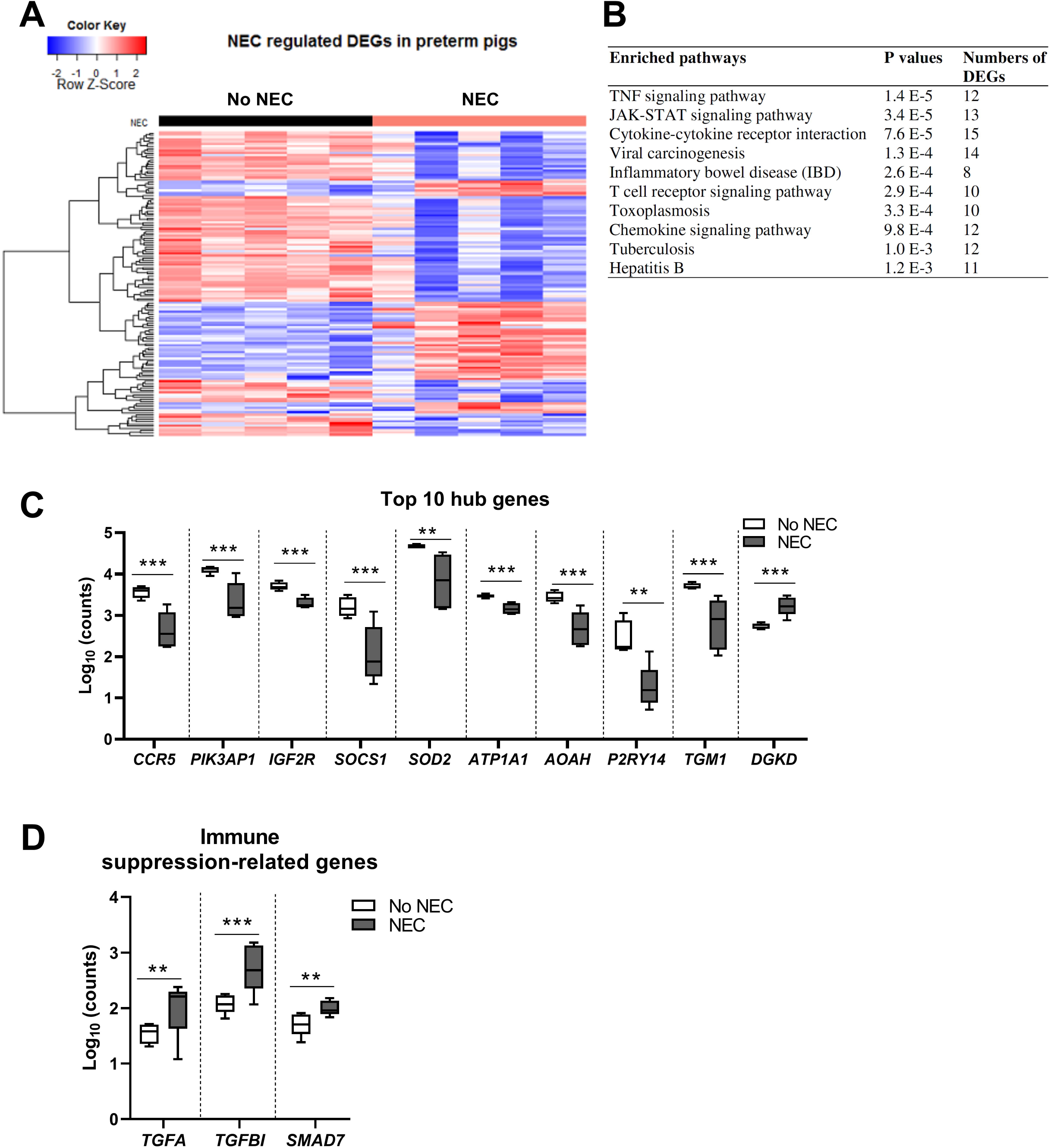
Blood transcriptomic profiles in pigs with and without NEC on d 9. Expression heatmap with differentially expressed genes (DEGs) between pigs with and without NEC (A). Top 20 enriched pathways from down-regulated genes in NEC pigs with FDR adjusted P values and number of genes involved in each enriched pathway (B). Expression levels of top 10 identified hub-genes among all DEGs between pigs without and with NEC (C) and genes related to immune suppression up-regulated in NEC pigs (D). n = 5 in each group of NEC or no NEC, with or without LPS stimulation. Values in D-E are means ± SEM. **, *** P < 0.01 and 0.001, respectively.

## Discussion

Gut bacterial translocation has recently been discussed as one of the main routes inducing bacteremia and sepsis^29,30^. In both septic preterm infants and animals, bacteria identified from positive blood cultures are often associated with the most abundant taxonomic groups in the gut microbiome^10,21,31^. Enteral administration of virulent bacterial strains also leads to bacterial translocation into the circulation and many systemic organs causing LOS^32^. The compromised and leaky gut barrier in NEC conditions may facilitate bacterial translocation and can be one of the reasons explaining that a fraction of NEC patients later experience one or more episodes of LOS^20,21^. Now with the current study, we provided evidence for NEC lesions programming the systemic immune system to a status of immune suppression, which may together with gut bacterial translocation contribute to the increased susceptibility to secondary infection and LOS in patients with antecedent NEC suspicion or diagnosis.

With multiple data collected at two different time points on d 5 and d 9, we were able to evaluate NEC incidence and severity during progression of the disease. Importantly, NEC incidence diagnosed by macroscopic lesions at euthanasia was identical on d 5 and d 9, suggesting that NEC found on d 9 had likely already occurred on d 5, and that the mucosal and systemic status on d 9 was a reflection of NEC effects. Strikingly, the severity of NEC lesions on d 9 was clearly less severe than on d 5, indicating the sub-clinical NEC lesions in some preterm pigs being resolved gradually. This important observation supports the fact that up to 70% of patients diagnosed with NEC recover over time without the needs of surgery^26,27^. It is possible that antibiotic treatment in some NEC cases associated with bacterial overgrowth may release the burden of high bacterial load in the gut, but the healing of NEC lesions may also occurs via specific host responses to avoid necrosis requiring surgery.

NEC lesions in preterm pigs here were characterized with decreased gut mucin-containing goblet cell density and increased blood neutrophil counts at early phase on d 5, followed by an infiltration of neutrophils and/or macrophages (MPO^+^ positive cells) and T lymphocytes (CD3^+^ lymphocytes) at later phase on d 9. These tendencies were even more significant in those pigs with more severe NEC lesions. These phenomena may reflect the innate immune defense from goblet cells releasing mucin as well as signaling from the gut inflammation priming granulopoiesis in early phase of NEC, followed by gut-homing immune cells at later phase on d 9. The influx of immune cells found in NEC tissues in our study were in line with many other NEC models in rodents^33,34^. Additionally, NEC occurrence during the first 9 days of life in our study was not associated with any changes in the composition of gut microbiota at either time points. This is consistent with previous reports showing no association between NEC occurring during the first month of life of very preterm infants and their gut microbiome compositions^10,28^. Importantly, changes in gut microbiome preceding NEC (increase in abundance of Gammaproteobacteria and decreased diversity) only occurred after the first month of life in preterm infants born at or before 27 weeks of gestation^28^. As gestational age at birth is disproportionally correlated with the age at NEC onset^35^, NEC occurrence during the first month of life, when bacterial colonization is extremely dynamic, may be more determined by the balance between the immune competence and bacterial overgrowth rather than by colonization of specific bacterial groups.

Many studies have characterized the changes of blood and gut cell subsets few days before or at the time of NEC diagnosis and showed increased numbers of inflammatory cells (gut CD4^+^ T effector memory cells, blood IL-17 producing Treg and CCR9^+^CD4^+^ T cells) or decreased numbers of immune suppressive cells subsets (gut Treg)^12,36–38^. The status at those NEC phases likely reflect the active immune responses of the host in an effort to resolve occurring gut inflammation. This is similar to the increased levels of inflammatory cytokines at the time close to diagnosis of LOS in preterm infants^16^. In contrast, the immune status following NEC diagnosis has not been focused. Now with the current study, we observed that preterm pigs with NEC on d 9, but not d 5, showed increased number of blood Treg, decreased frequency of phagocytic neutrophils as well as impairment of LPS-induced cytokine secretion and mRNA responses for genes related to innate and adaptive immunity in whole blood. The transcriptomic profile of NEC pigs on d 9 also demonstrated clear depression of pathways related to innate immune signaling and Th1 cell polarization, including TNF, STAT and T cell receptor signaling^39,40^. Collectively, our data indicate that NEC lesions gradually programmed the systemic immune system in preterm pigs to a state of immune suppression. The active inflammatory responses at earlier phases of NEC may consume most of the stored energy in the circulating immune cells, resulting in an exhausted and paralytic state at a later phase^7^. We postulate that NEC-induced systemic immune suppression may be an important contributing factor interacting with bacterial translocation across the compromised gut barrier to predispose NEC patients to increased risks of secondary infection and LOS^20,21^. The characteristics and mechanisms of immunosuppression induced by NEC may share some similarities to that of immunosuppressive status found in late stages of septic infants and elderly patients, which also predisposes those patients to secondary infection episodes and organ failures^16,17^.

It is also noteworthy that a series of genes from the transcriptomic profile related to glycolysis and TCA cycle (*PDHX, PDK3, ADPGK, PCK2*), oxidative phosphorylation and ATP synthesis (*CYP1B1* and genes involved in ATPase activities) consistently showed lower levels in pigs with vs. without NEC. This supports our postulation of NEC-induced energy deprivation in circulating immune cells with down-regulated pathways synthesizing ATP and decreased immune competence. Similar phenomena of impaired immunity and energy production-related pathways were observed in monocytes of preterm vs. term infants, as well as whole blood of preterm pigs born with vs. without prenatal inflammation^9,24^. It would be important to further elucidate the responses of NEC pigs to *in vivo* infection challenges in future studies to characterize in details their programmed immunometabolic status and confirm their increased susceptibility to secondary infection and sepsis.

In conclusion, we demonstrated clear patterns of systemic immune suppression following NEC lesions in newborn preterm pigs. The effects were observed in animals with both mild and severe sub-clinical lesions. The current study provided important evidence for the compromised immune status in NEC patients, which may explain mechanisms of impaired immune defense to secondary infectious challenges and a proportion of NEC patients later developing one or more episodes of LOS. Finally, it is noteworthy that NEC occurrence in this study was mainly sub-clinical without manifestation into severe clinical symptoms, thereby reflecting a state of NEC suspicion or medical NEC in preterm infants. Still, our data suggest careful management of preterm infants with mild signs of gut dysfunction to avoid secondary infection due to the negative impact on the systemic immune system.

## Materials and Methods

### Preterm pig experimental procedures and sample collection

All animal procedures were approved by the Danish National Committee of Experimentation. A preterm pig cohort was set up with 113 pigs (Landrace x Yorkshire x Duroc) delivered by caesarean section at day 106 of gestation (~90%, term at d 117) from ten sows. After delivery, pigs were transferred to individual incubators with supplemental oxygen (0.5-2 L/min) for the first 24 h. Each pig was inserted a vascular catheter (4F, Portex, Kent, UK) via the umbilical artery for parenteral nutrition (Kabiven, Fresenius Kabi, Uppsala, Sweden) and blood sampling, and an orogastric catheter (6F, Portex) for enteral nutrition with milk diets (increasing amount of 16-112 ml/kg/day of bovine colostrum or bovine milk-based formula, 3300-3500 KJ/L) until postnatal day (d) 9. To provide passive immunity, each pig received 16 mL/kg maternal plasma via the umbilical catheter during the first 24 h of life. Pigs were continuously monitor in individual incubator. On d 5, 36 pigs from several litters were planned for euthanasia for gut tissue collection and macroscopic NEC diagnosis. The remaining pigs were reared until euthanasia at postnatal d 9. When severe NEC clinical symptoms appeared before the planned euthanasia, pigs were also euthanized according to humane endpoints. At euthanasia, gut tissues were dissected, and the small intestine was equally divided into three regions: proximal, middle and distal small intestine. Macroscopic lesions were scored based on a previously documented scoring system from 1 to 6, based on the degree and severity of hyperemia, edema, hemorrhage, pneumatosis and necrosis^41^. A pig with a score of three or higher in any of the small intestinal regions or colon was designated as NEC. A pig with the highest regional score of 3-4 and 5-6 was stratified as mild and severe NEC, respectively. Small intestinal and colonic tissues were snap-frozen or fixed in paraformaldehyde 4% for later analysis. Fixed tissues were embedded in paraffin and 5 μm sections were used for histology.

On d 5 and 9, blood samples were collected for all pigs from the arterial catheter. Fresh blood was used for hematology by an automatic cell counter (advia 2120i Hematology System, Siemens, Germany), flow cytometry, and *ex vivo* stimulation with LPS. Blood (200 μl) were also mixed with 520 μl mixture of lysis/binding solution concentrate and isopropanol (MagMax 96 blood RNA isolation kit, Thermofisher, Roskilde, Denmark), and stored at −80°C until RNA extraction for transcriptomics and qPCR analysis.

### *In vivo* test and gut tissue analysis

Precisely three hours prior to the planned euthanasia, a gut permeability test was performed by enteral administration of a solution containing 5% (w/v) of lactulose and mannitol and urinary levels of these two molecules were measured at euthanasia as previously described^41^. Villus height and crypt depth in the three small intestinal regions were measured on images from H&E stained fixed tissues using Image J (National Institutes of Health, Bethesda, MD, USA)^41^. Colonic and distal small intestinal mucin-containing goblet cell density was determined following Alcian Blue and Periodic acid-Shiff staining as previously described^42^. Colon and distal small intestinal tissues were also stained for CD3 (T cell marker) and MPO (neutrophil/macrophage marker) for indication of inflammatory cell infiltration, using anti-porcine CD3 (Southern Biotech, Birmingham, AL, USA) and anti-human MPO (Agilent, Glostrup, Denmark). CD3 was quantified by positive cell area fraction using Image J, whereas MPO was graded from 1-7 according to cellular density (low, medium, high) and degree of tissue inflammation (none, focal, multifocal, diffuse, ulceration) by a blinded investigator. Cytokines (IL-8, IL-6, TNF-α and IL-1β) in the frozen distal small intestine and colon were measured by porcine specific ELISA assays (R&D Systems, Abingdon, UK).

### Gut microbiome

Enema samples (d 5) or colon content (d 9) of the same selected pigs reared to postnatal d 9 from three litters were used for total DNA extraction (PowerSoil DNA isolation kit, MoBio Laboratories). Thereafter, 16S rRNA gene (V3 region) amplicon sequencing (Illumina, San Diego, CA, USA) and downstream bioinformatics were performed as previously described^43^. Raw reads were analyzed and zero-radius operational taxonomic units (zOTUs) were constructed using UNOISE algorithm^44^. Sample counts were rarified to 4800 for calculation of alpha and beta diversity, and the unweighted and weighted Unifrac distance metrics were visualized by principal coordinate analysis (PCoA).

### Whole transcriptome shotgun sequencing

Stored blood samples collected on d 9 from selected 8 NEC and 8 healthy pigs across 3 litters were used for RNA extraction using MagMax 96 blood RNA isolation kit (Thermofisher) for transcriptomics by whole transcriptome shotgun sequencing as previously described^24^. Briefly, sequencing libraries were constructed with NEBNext Ultra RNA Library Prep Kit for Illumina (New England Biolabs, Ipswich, MA, USA) and libraries were sequenced using Illumina Hiseq 4000 platform (Illumina). After that, 150 bp paired end raw reads were trimmed and then were aligned to the Sscrofa 11.1 genome using TopHat^45^, and gene information was obtained from Ensembl (www.ensembl.org/Sus_scrofa/Info/Index). Gene counts were generated and differentially expressed genes (DEGs) between pigs with vs. without NEC with cutoff q values < 0.05 using DESeq2^46^. Lists of DEGs with mean expression levels and adjusted P values were listed in Supplementary Table S1. Biological process and KEGG pathway enrichment was performed using DAVID Bioinformatics Resources, with cutoff of adjusted P values < 0.05 for enriched pathways (Supplementary Table S2).

### Immune assays with whole blood LPS stimulation, neutrophil phagocytosis and cell profiling

Fresh blood samples on d 5 and 9 (200 μl) was stimulated with LPS (1 μg/mL, O127:B8 from *E.coli*, SigmaAldrich) for 5 h at 37°C and 5% CO_2_. After stimulation, a fraction of blood was mixed with the lysis/binding solution prepared in isopropanol (Thermofisher) and stored at −80°C for qPCR. The other fraction of blood was centrifuged (2000 x g, 10 min, 4°C) and plasma was analyzed for TNF-α and IL-10 (porcine specific ELISA kit, R&D Systems, Abingdon, UK). For qPCR, blood RNA was extracted and converted into cDNA ^47,48^, and qPCR for a panel of selected 23 genes related to innate, adaptive immunity and cellular metabolism was performed using QuantiTect SYBR Green PCR kit (Qiagen) on LightCycler 480 (Roche). Relative gene expression was normalized to the housekeeping gene *HPRT1*. Primer-Blast (http://ncbi.nlm.nih.gov/tools/primer-blast) were used to designed primers and their sequences were described in Supplemental Table S3.

Fresh blood samples on d 5 and 9 were also analyzed for neutrophil phagocytosis function and T cell subset profiling. Phagocytosis assay was performed with 100 μl blood using the pHrodo Red E.coli (560/585 nm) Bioparticles Phagocytosis Kit for Flow cytometry (Thermofisher, Roskilde, Denmark) and the Accuri C6 flow cytometer^43^. pHrodo^+^ neutrophils were identified as neutrophils exerting phagocytosis while median fluorescence intensity of pHrodo^+^ neutrophils demonstrated phagocytic capacity (number of bacteria being engulfed). T cell subset profiling was performed flow cytometry^24,49^. Following erythrocyte lysis (10 x BD FACS Lysing solution diluted with sterile water, BD Biosciences, Lyngby, Denmark), leukocyte permeabilization (Fixation/Permeabilization buffer, for 30 min at 4°C in the dark, washed twice by permeabilization buffer, all from Thermofisher), Fc receptor blocking (porcine serum for 15 min at 4°C in the dark), leukocytes were stained with a mixture of 4 antibodies: PerCP-Cy5.5 conjugated anti-pig CD3 antibody, FITC-conjugated anti-pig CD4 antibody, PE-conjugated anti-pig CD8 antibody (all three from Biorad, Copenhagen, Denmark), and APC-conjugated anti-mouse/rat Foxp3 antibody (Thermofisher). Corresponding negative controls were used as isotype controls. Stained cells were analyzed by Accuri C6 flow cytometer (BD Biosciences). The following T cell subsets were identified: helper T cells (Th, CD3^+^CD4^+^CD8^-^ lymphocytes), cytotoxic T cells (Tc, CD3^+^CD4^-^ CD8^+^ lymphocytes) and regulatory T cells (Treg, CD3^+^CD4^+^Foxp3^+^ lymphocytes).

### Statistics

All statistics were performed using R, version 3.4.3. Categorical data (NEC incidence) were analyzed using Fisher’s exact test. Continuous data (except transcriptomic and gut microbiome data) were analyzed by linear models with NEC (no NEC vs. NEC; or no NEC vs. mild NEC vs. severe NEC) and litters as fixed effects using lmer functions. Post-hoc Tukey tests were used when comparing groups with different NEC severities. Data were log transformed if necessary. Residuals and fitted values were evaluated for normal distribution and variance homogeneity. P values < 0.05 were regarded as statistical significance and P values in the range of 0.05-0.1 were considered as tendencies of being significant. For transcriptomic data, heatmaps were generated by heatmap. 3 function. To identify hub-genes among DEGs between pigs with vs. without NEC, all DEGs were analyzed for correlation (Spearman) and significant correlations (Spearman’s Rho >0.8 and adjusted P-value < 0.05) were further analyzed for degree of interactions and betweenness centrality (Cytoscape).

## Author contributions

DNN designed the study. DNN, SR, YH and XP carried out the experiments and laboratory analysis. SR, YH and XP performed data analysis. SR and DNN wrote the manuscript. SR, YH, FG, DN, PTS and DNN provided critical interpretation and revised the manuscript. SR and DNN had primary responsibility for the final content and all authors approved the final paper.

## Acknowledgement

The authors would like to thank Thomas Thymann, Yanqi Li, Anders Brunse, Elin Skytte, Jan C. Povlsen, Kristina Møller and Brita Karlsson for the assistance in pig experiments and analyses. The study was supported from the STIMMUNE and NEOCOL projects granted by the Arla Food for Health (DNN and PTS) and Innovation Foundation Denmark (PTS), respectively. The authors declare no conflicts of interest.

